# Combinatorial inhibition of LSD1 and Menin induces therapeutic differentiation in AML

**DOI:** 10.1101/2025.11.09.687496

**Authors:** María F. Carrera Rodríguez, Joshua Rico, Manaswini Vijayaraghavan, Fangxue Yan, Austin King, Ricardo Petroni, Nicolae Adrian Leu, Haley Goodrow, Kathrin Bernt, Gerald McGeehan, M. Andrés Blanco

## Abstract

Acute myeloid leukemia (AML) is characterized by differentiation arrest and uncontrolled proliferation. Differentiation therapy aims to treat AML by de-repressing latent myeloid maturation programs to induce cell cycle arrest and subsequent cell death. This approach is curative in the promyelocytic AML subtype, but has met with limited success in other subtypes. Genes such as LSD1 have emerged as intriguing non-APL AML differentiation therapy targets, but results as monoagents in clinical trials have been mixed. Here, we performed differentiation-specific CRISPR screens to identify targets whose inhibition synergizes with LSD1 inhibition to induce terminal differentiation of non-APL AML cells. Intriguingly, the MLL co-factor Menin scored as the top hit. Using cell lines, primary patient samples, and mouse AML models, we find that dual inhibition of LSD1 and Menin is a highly promising approach for differentiation therapy. Mechanistically, we determine that inhibition of Menin downregulates drivers of proliferation and stemness such as MEIS1, and inhibition of LSD1 induces inflammatory and interferon-related pro-myeloid differentiation expression programs. Surprisingly, we find that this combination is effective in selected AML models without mutations in MLL or NPM1, thus nominating dual inhibition of LSD1 and Menin as an attractive therapeutic approach for a mutationally diverse set of non-APL AMLs.

**Highlights:**

- Inhibition of LSD1 and Menin synergizes to induce differentiation of MLL-r and MLL-WT AMLs.
- Inhibition of Menin downregulates drivers of proliferation and stemness.
- Inhibition of LSD1 induces differentiation-associated inflammatory and interferon responses.
- LSD1 and Menin occupy different areas of the genome.

## Introduction

Acute myeloid leukemia (AML) is a devastating disease that accounts for roughly 20,000 deaths in the United States annually^1^. Treatment options have not changed for most patients for several decades, as intensive chemotherapy typically remains the standard of care. As the majority of diagnosed AML patients will die from their disease, novel treatments are of great need.

While AMLs are mutationally heterogenous^2^, all feature a prominent differentiation block that prevents them from maturing, exiting the cell cycle, and dying^3^. A breakthrough in AML oncology came with the discovery that the differentiation block could be targeted. The combination of ATRA and ATO degrades the PML-RARα fusion protein that drives acute promyelocytic leukemia (APL), leading to cell cycle arrest and cell death with minimal toxicity^4^. This treatment approach, known as differentiation therapy, is the standard of care for APL patients and is 95% curative^5^. However, while differentiation therapy has been transformative for APL, it has met with limited success in non-APL AMLs – which represent 85-90% of AML overall.

Recent findings suggest that targeting chromatin regulators may offer a promising avenue for differentiation therapy of non-APL AMLs^6^. For example, IDH inhibitors act largely through inducing differentiation in *IDH1/2-*mutant AML and have been clinically successful^7^. Another attractive target is the H3K4me1/2 demethylase LSD1. Several studies have shown that LSD1 is a prominent driver of the non-APL AML differentiation block, and that LSD1 inhibitors (LSD1i) induce varying degrees of differentiation in a variety of AML mutational subtypes^8–10^. However, LSD1i efficacy in the clinic has been limited due to associated toxicity^11^. This suggests that LSD1i may be most effective at a lower dose in combination with other inhibitors. Along these lines, recent studies have proposed LSD1i combination therapy partners. 5-Azacytidine has been shown to combine effectively with LSD1i in the treatment of *TET2-*mutant AML^12^. Very recently, inhibition of the β-Catenin kinase GSK3 was also shown to combine with LSD1 to induce dramatic differentiation in a variety of AML subtypes^13^. However, to date no studies have reported chromatin regulators whose inhibition synergizes with LSD1i to induce non-APL AML differentiation.

Here, we leverage a genetic screening strategy to find that inhibition of the MLL co-factor Menin synergizes with LSD1i to induce therapeutic differentiation in non-APL AML. Surprisingly, this combination was found to be effective not only in AMLs harboring MLL rearrangements, but also in AMLs lacking mutations in MLL or NPM1. This runs counter to the prevailing thinking that Menin inhibitors will only be effective in MLL-rearranged and NPM1-mutant AMLs. Accordingly, our study reports a novel, epigenetic-based drug combination that may be effective for a wider variety of AMLs than would have previously been predicted.

## Results

### Differentiation-focused LSD1 synergy screen identifies Menin as the top hit

To identify potential LSD1 inhibitor (LSD1i) combination therapy partners, we performed an LSD1i “synergy screen.” We used the ER-Hoxb8 differentiation arrest model^14^, as it provides a genetically-defined, primary cell context that has been successfully used for prior AML screens^15^. ER-Hoxb8 cells are MLL-WT but driven by similar Hox-like gene expression programs as MLL-r AML. We delivered a chromatin-focused CRISPR sgRNA library to Cas9-ER-Hoxb8 cells and then split the cells into a DMSO-treated branch and a branch treated with the LSD1 inhibitor ORY-1001 at 10 nM. Cells were cultured for one week and then both branches were sorted based on the myeloid differentiation-associated surface marker CD11b (Figure S1). We collected the top 2% (highly differentiated) and bottom 2% (lowly differentiated) CD11b cells and used next generation sequencing to identify sgRNAs enriched in the CD11b high population in each screen branch. These sgRNAs represent genes whose inhibition induces differentiation. The screen was technically sound, achieving high coverage and expected levels of variance in sgRNA abundances in CD11b high and CD11b low populations in both screen branches (Figure 1A). The screen was also biologically successful, with the DMSO-treated branch capturing as top hits several known drivers of AML differentiation arrest, such as Mediator complex subunits, Dot1l, Kat6a, and Rcor1, the structural subunit of the LSD1-containing co-REST complex.

**Figure 1.**
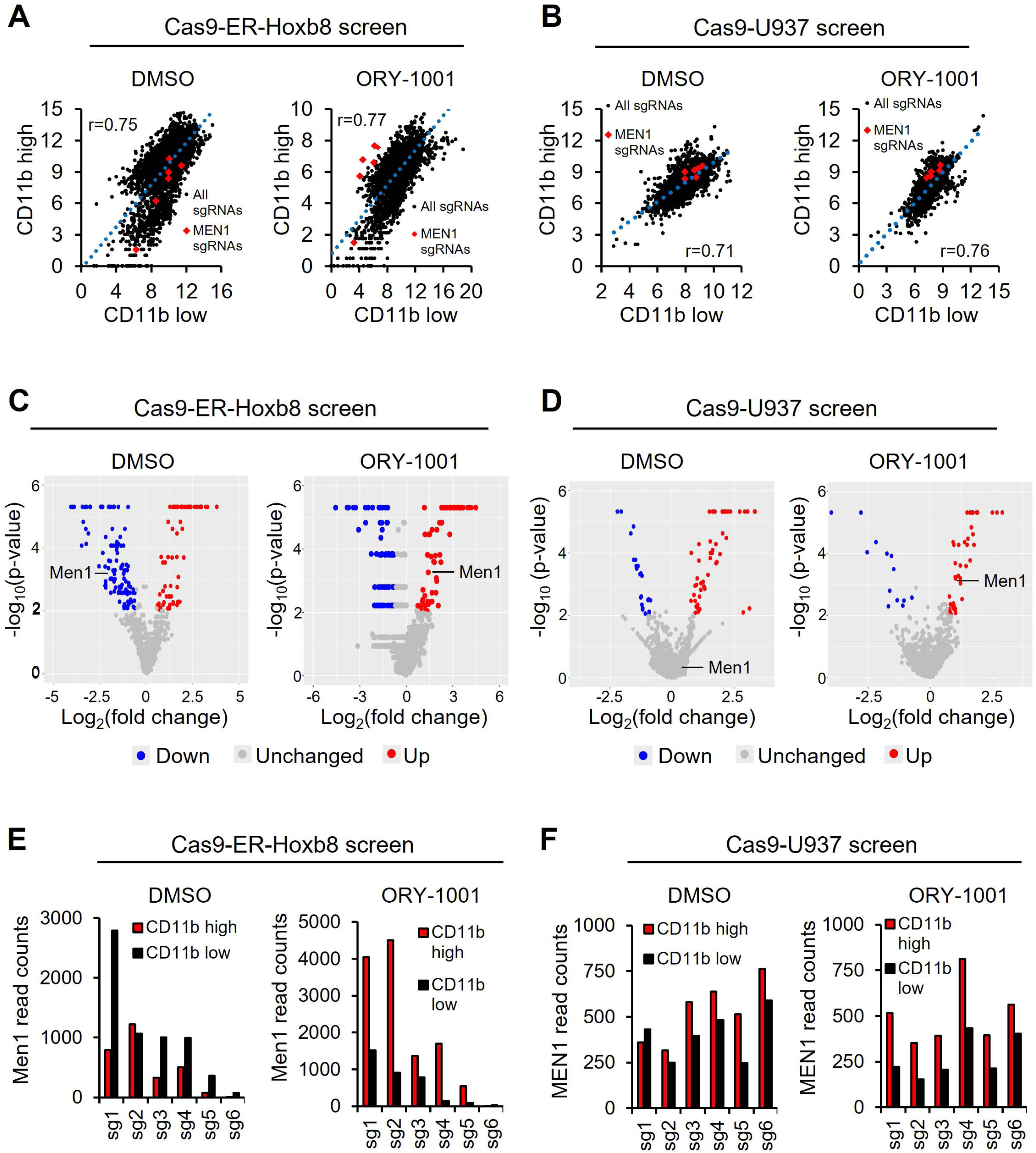
Screen identifies Menin as a potential LSD1 combination differentiation therapy partner. **A.** Correlation in read counts between CD11b low (lowly differentiated) and CD11b high (highly differentiated) cell populations from gain-of-differentiation screen performed in Cas9-ER-Hoxb8 cells. **B.** Correlation in read counts between CD11b low (lowly differentiated) and CD11b high (highly differentiated) cell populations from gain-of-differentiation screen performed in Cas9-U937 cells. **C.** Volcano plots showing Cas9-ER-Hoxb8 screen hits enriched in CD11b high cells (red), CD11b low cells (blue), and non-enriched (grey). **D.** Volcano plots showing Cas9-U937 screen hits enriched in CD11b high cells (red), CD11b low cells (blue), and non-enriched (grey). **E.** Read counts of Menin in CD11b high and low populations from Cas9-ER-Hoxb8 screen. **F.** Read counts of Menin in CD11b high and low populations from Cas9-U937 screen.

We then looked for genes that scored significantly in the ORY-1001-treated branch but not the DMSO-treated branch, as these represent genes whose inhibition may synergize with ORY-1001 to induce differentiation. Intriguingly, there was only one such gene in this library of >800 genes: the MLL co-factor Menin (Figures 1C and 1E). We then performed an analogous screen in the MLL-WT human AML cell line U937. This screen was also technically and biologically successful (Figure 1B). Remarkably, the same result was obtained – Menin was the only hit that scored in the ORY-1001-treated branch but not the DMSO-treated branch (Figures 1D and 1F). This suggests that combinatorial inhibition of LSD1 and Menin may synergistically induce differentiation in AML cells. This is especially surprising given that Menin inhibition is thought to only be effective in MLL-r or NPM1 mutant-driven AMLs.

We sought to confirm screen results. As there is a highly effective Menin inhibitor, SNDX-5613^16^, we used small molecule inhibition, rather than genetic KO, of Menin for the remainder of the study. In ER-Hoxa9 cells, which have a built-in Lysozyme-GFP differentiation reporter, Menin inhibition via SNDX-5613 had almost no effect on GFP levels up to 50 uM. Similarly, LSD1 inhibition via ORY-1001 had minimal effect on GFP up to 10 nM. However, combinatorial inhibition markedly induced the GFP differentiation reporter starting at 1 uM SNDX-5613 and as low as 2.5 nM ORY-1001 (Figure 2A). As differentiation should lead to proliferation arrest, we next tested the effect of the ORY-1001+SNDX-5613 combination (hereafter, “c ombo”) on proliferation/viability via CellTiter-Glo assays. While SNDX-5613 used alone had an IC50 of 0.47 uM, addition of 5 nM ORY-1001 markedly reduced the IC50 to 0.20 uM (Figure 2B). These effects produced extremely high ZIP synergy scores, reaching the upper 40s (Figure 2C). Importantly, we showed that these results are not unique to the ORY-1001 inhibitor. We found that 100 nM of the GSK-LSD1 inhibitor also synergized with SNDX-5613 to induce CD11b in MLL-AF9 cells (Figure S2A).

**Figure 2.**
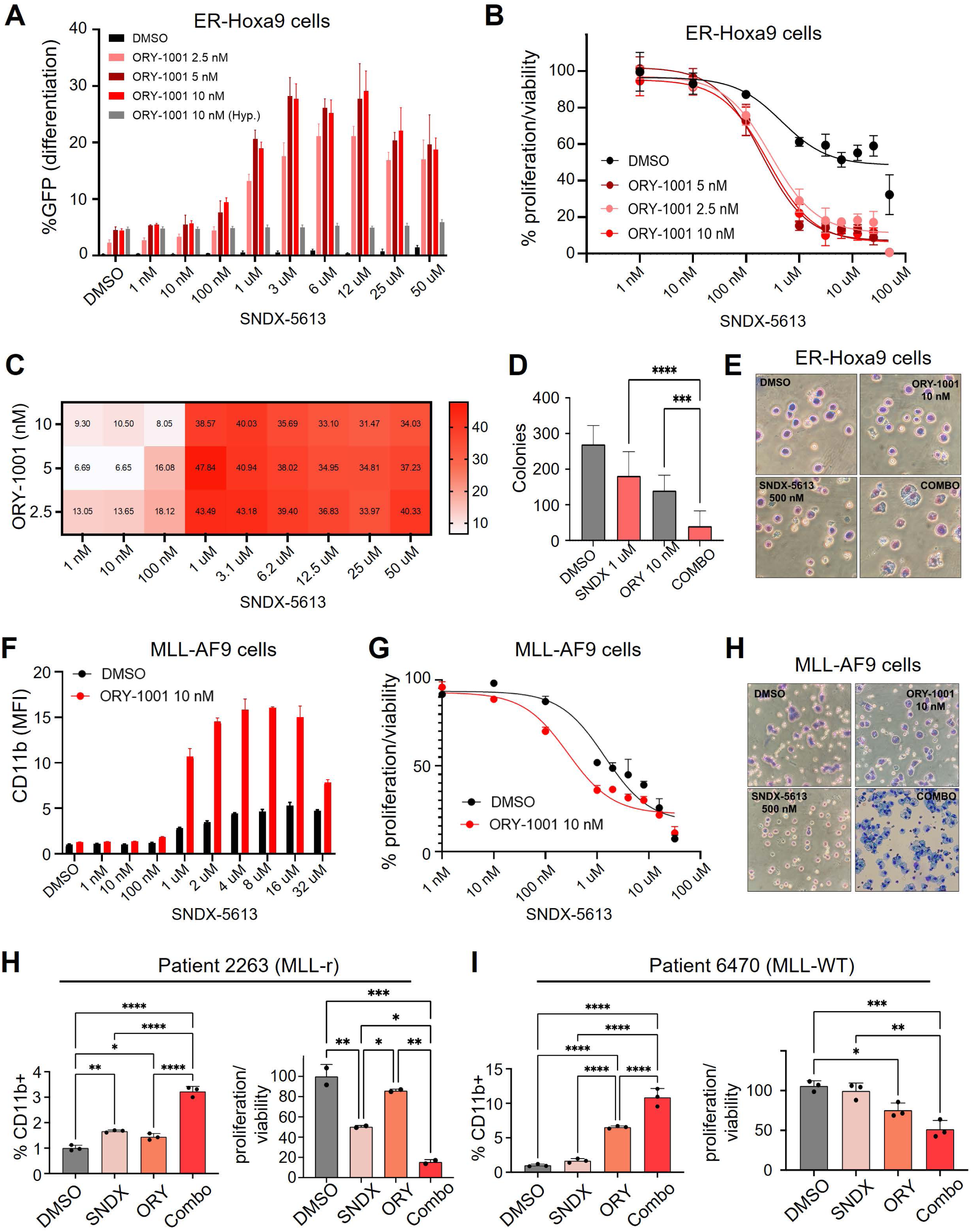
Combinatorial inhibition of LSD1 and Menin induces differentiation in AML cells. **A.** Differentiation response of ER-Hoxa9 cells treated with SNDX-5613, ORY-1001, both, or neither. GFP is driven by internal Lysozyme-GFP differentiation reporter built in to ER-Hoxa9 cells. **B.** CellTiter-Glo assays for proliferation/viability responses of ER-Hoxa9 cells treated with SNDX-5613, ORY-1001, both, or neither. Grey bars indicate hypothetical additive of effect of ORY-1001 alone combined with SNDX-5613 alone at the given concentrations. **C.** SynergyFinder synergy scores for ORY-1001+SNDX-5613 combination based on CellTiter-Glo data. **D.** Colony formation assay in ER-Hoxa9 cells after drug treatments. **E.** Hema3 staining of ER-Hoxa9 cels after drug treatments. **F.** Differentiation response of MLL-AF9 cells treated with SNDX-5613, ORY-1001, both, or neither using CD11b as a readout for differentiation. **G.** CellTiter-Glo assays for proliferation/viability responses of MLL-AF9 cells treated with SNDX-5613, ORY-1001, both, or neither. **H.** CD11b (left) and CellTiter-Glo (right) responses to drug treatments in patient 2263’s AML cells cultured *ex vivo.* **I.** CD11b (left) and CellTiter-Glo (right) responses to drug treatments in patient 6470’s AML cells cultured *ex vivo*.

If the ORY-1001+SNDX-5613 drug combination works via inducing differentiation, we would expect self-renewing ability to be lost. Indeed, while SNDX-5613 and ORY-1001 reduced colony formation by roughly two-fold when used alone, their combination almost completely ablated this activity (Figure 2D). Finally, we evaluated the effects of the combo on cell morphology and histology using Hema-3 staining. We found that most cells looked moderately to strongly differentiated, with some appearing terminally (fully) differentiated (Figure 2E). These findings were encouraging, especially given that ER-Hoxa9 is an extremely conservative model of differentiation arrest, with it being very challenging to achieve terminal differentiation due to the continued high activity of the Hoxa9 transcription factor regardless of cell treatment.

As ER-Hoxa9 cells are MLL-WT, we investigated the effects of the combo in MLL-r AML using primary mouse AML cells transformed by retroviral overexpression of MLL-AF9. Results were largely similar (Figures 2F-2H), except that, as expected, SNDX-5613 used alone produced a greater differentiation response than it did in ER-Hoxa9 cells. Notably, almost 100% of MLL-AF9 cells treated with the combo appeared terminally differentiated (Figure 2H). As an additional test for differentiation, we performed superoxide anion (SOA) assays, which test for the ability of cells to produce the SOA free radical in response to a stimulus that mimics an infection. This activity characterizes mature myeloid cells such as neutrophils and macrophages. Intriguingly, while control and monoagent-treated cells were unable to mount a free radical response, combo-treated cells responded with robust SOA production in response to PMA stimulus (Figure S2B). We next tested the combo in primary AML patient samples cultured *ex vivo.* While responses varied across samples tested, we identified both MLL-r and MLLL-WT/NPM1 WT samples in which the combo strongly potentiated CD11b levels and reduced proliferation (Figures 2H and 2I). These results indicate that combo efficacy is not limited to the ER-Hoxa9/b8 model and is not an artifact of long-term *in vitro* cell culture.

We further investigated the effect of the drug combo on AMLs that were driven by MLL-r, NPM1c mutation, or neither. We performed proliferation/viability and CD11b response assays in three MLL-r cell lines (MOLM13, MV4;11, and THP1), three MLL-WT cell lines (U937, HL60, and the chronic myeloid leukemia line (CML) K562), and one NPM1c cell line (Oci-AML3). Remarkably, though responses varied according to dose, there was at least one concentration in which the drug combo both induced CD11b and reduced proliferation/viability synergistically or near-synergistically in all cell lines tested (Figures 3A and 3B). There were cases in which higher SNDX-5613 concentrations induced growth/proliferation inhibition on its own; however, synergy was still observed at a lower SNDX-5613 concentration in these cases. These results confirm that AMLs of a wide variety of genetic backgrounds, including MLL-WT and NPM1 WT, can respond strongly to the SNDX-5613/ORY-1001 drug combo.

**Figure 3.**
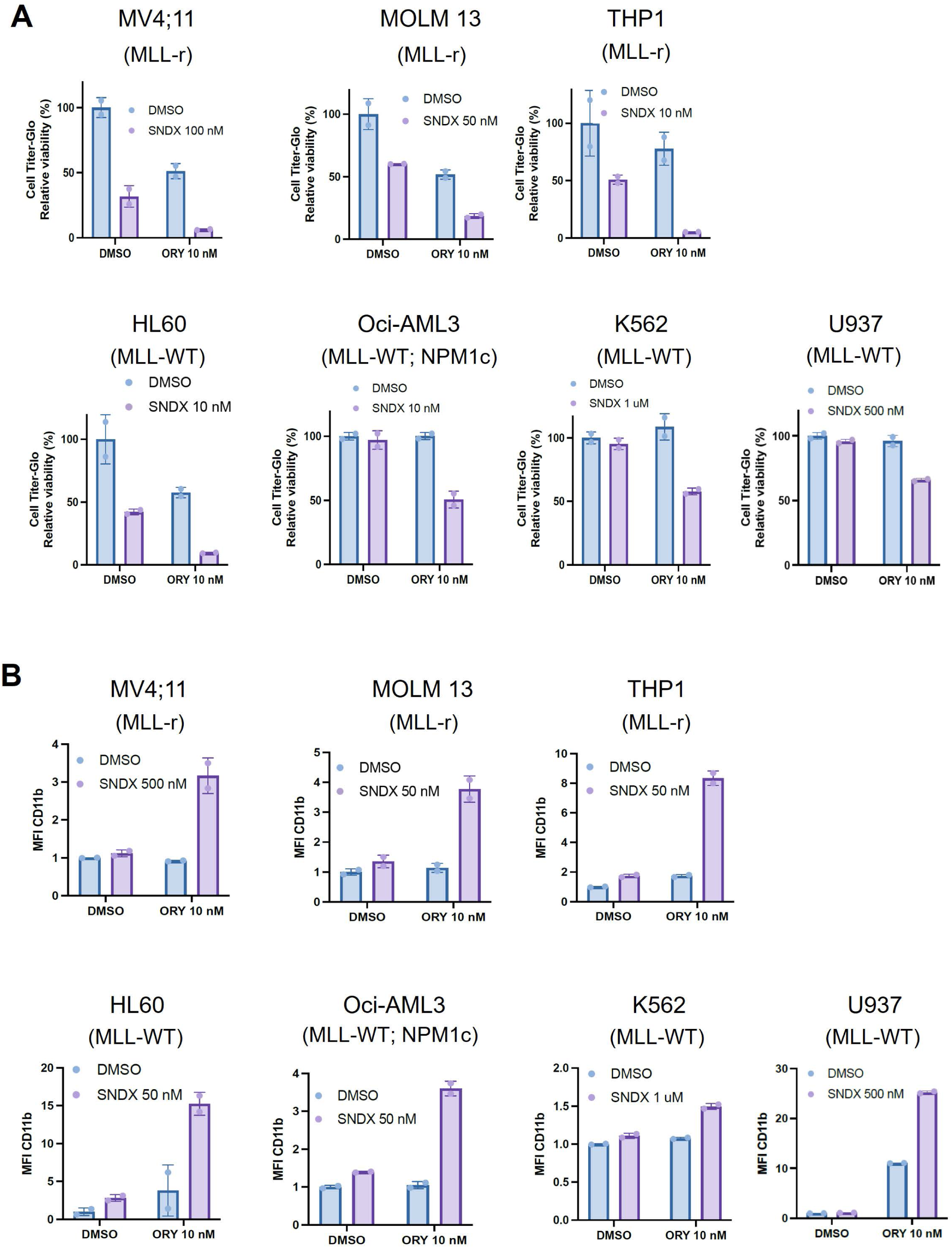
Combinatorial inhibition of LSD1 and Menin induces differentiation in a panel of AML cell lines. **A.** CellTiter-Glo assays of AML cell lines after treatments with ORY-1001, SNDX-5613, the combination, or neither. **B.** Differentiation response of cell line panel after treatment with SNDX-5613, ORY-1001, both, or neither using CD11b as a readout for differentiation.

We next considered the mechanisms by which inhibition of Menin and LSD1 induces differentiation. We performed RNA-seq on ER-Hoxa9 cells treated with 10 nM ORY-1001 (alone), 1 uM SNDX-5613 (alone), the combination, or DMSO after 24 or 96 hours of treatment. PCA plots were informative. At day 4, ORY-1001 alone produced a marked shift in PC1 but little change in PC2. SNDX-5613 induced the opposite trend; strong movement on PC2, but almost no change on PC1. Combo treatment appeared to combine these effects, with dramatic and roughly even movement on both PC1 and PC2 (Figure 4A). This result suggests that, interestingly, SNDX-5613 and ORY-1001 may be inducing different gene expression programs. This is contrary to most cases of synergy, in which each treatment acts on the same module, with both treatments being necessary to disable (or induce) the module.

**Figure 4.**
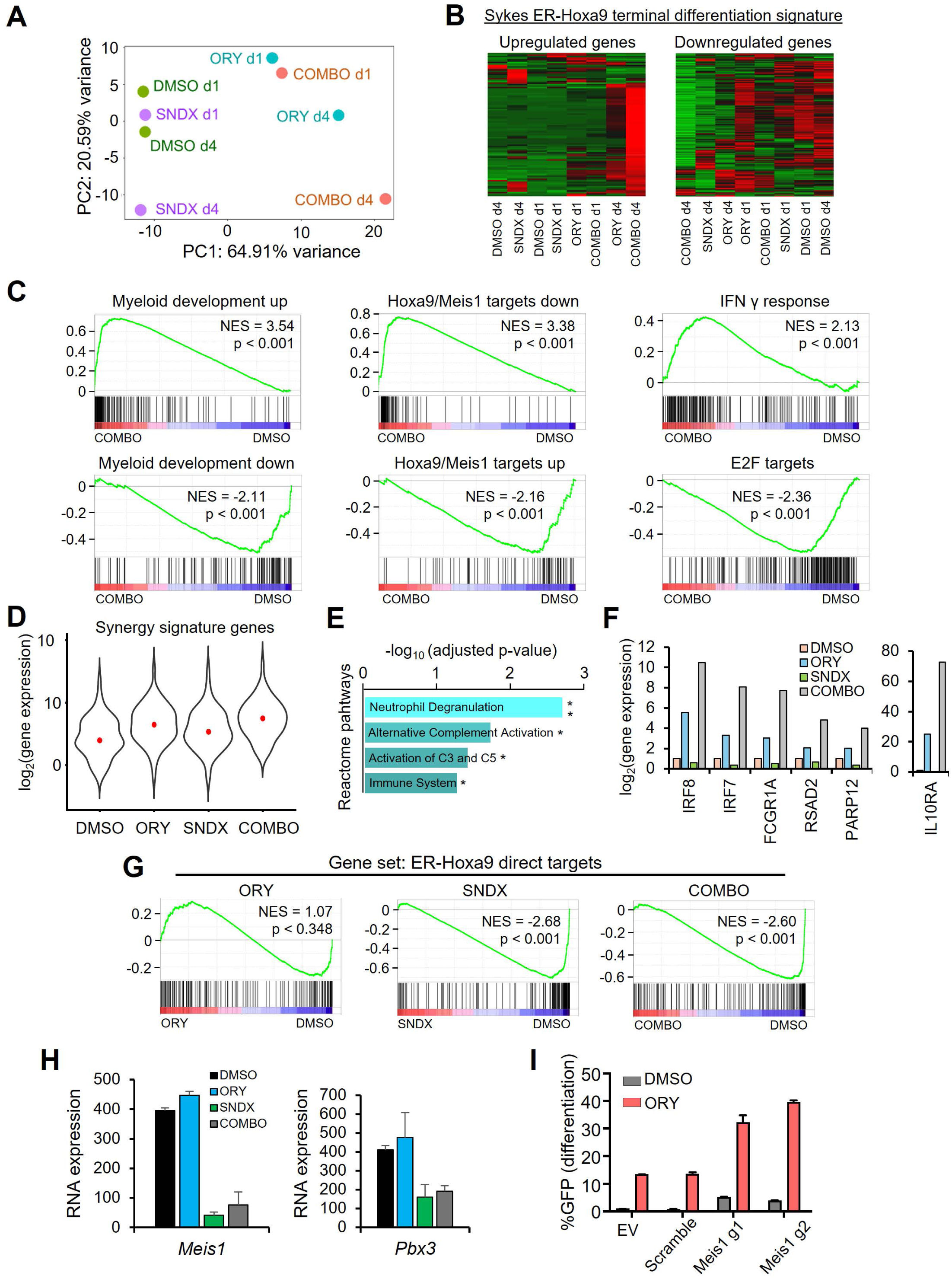
Analysis of transcriptional response of AML cells to drug combination treatment. **A.** PCA plot from RNA-seq of ER-Hoxa9 cells treated with ORY-1001, SNDX-5613, the combination, or neither for 1 or 4 days. **B.** Hierarchical clustering of drug-treated ER-Hoxa9 cells using the upregulated and downregulated components of the Sykes terminal myeloid differentiation signature. **C.** GSEA plots of RNA-seq from ER-Hoxa9 cells following drug treatments. **D.** Expression patterns of genes that are synergistically upregulated in ER-Hoxa9 cells treated with ORY-1001+SNDX-5613 drug combination. **E.** Reactome pathways enriched in genes synergistically upregulated in ER-Hoxa9 cells treated with the ORY-1001+SNDX-5613 drug combination. **F.** Expression patterns of selected interferon-related genes upon treatment with the ORY-1001+SNDX-5613 drug combination. **G.** GSEA plot showing enrichments of the set of mapped direct targets of Hoxa9 in RNA-seq of ER-Hoxa9 cells following drug treatments. **H.** RNA-seq expression of *Meis1* and *Pbx3* in ER-Hoxa9 cells following drug treatments. **I.** Induction of GFP differentiation reporter in ER-Hoxa9 cells upon knockout of *Meis1,* treatment with ORY-1001, both, or neither.

We then analyzed the gene expression programs induced by ORY-1001 and SNDX-5613. Clustering on the Sykes terminal myeloid differentiation signature (derived from ER-Hoxa9 cells^15^) indicated that combo treatment robustly upregulated genes that promote differentiation and downregulated genes that oppose differentiation (Figure 4B). Notably, ORY-1001 treatment (alone) clustered next to combo treatment in the upregulated genes, and SNDX-5613 treatment (alone) clustered next to combo treatment on the downregulated genes (Figure 4B). This suggests a potential model in which the most direct effects of LSD1 inhibition are de-repression of pro-myeloid differentiation genes and the most direct effects of Menin inhibition are downregulation of pro-stemness/proliferation genes.

We next took an unbiased approach to evaluating the effects of the drug combo by running the RNA-seq data through the GSEA database. In line with our findings thus far, the top gene set from the c2 chemical and genetic perturbations database in combo-treated cells was the Brown myeloid differentiation up signature (Figure 4C). The second most enriched gene set (of a total of 2,444 tested) was the set of genes downregulated by Hoxa9 and Meis1 expression, suggesting that Hoxa9/Meis1 repression targets are de-repressed upon combo treatment. Correspondingly, both the Brown myeloid differentiation down and Hoxa9/Meis1 up targets were markedly downregulated in combo treatment (Figure 4C). Among other strongly regulated gene sets, interferon signatures such as interferon Y were highly upregulated upon combo treatment, while E2F target genes were markedly downregulated (Figure 4C). The former result has been observed before in LSD1i-based differentiation-inducing drug combinations^13^, while the latter result presumably reflects the halted proliferation that characterizes terminal myeloid differentiation. Of note, all of these gene sets were also significantly induced/repressed in each monoagent treatment, just to markedly reduced extents in comparison to combo treatment.

We next investigated the set of genes that are induced by the combo but not by either alone. In this “synergy signature” (Figure 4D), the most enriched gene set in the Reactome database was the highly specific neutrophil degranulation gene signature, which characterizes the most advanced stages of neutrophil maturation (4E). Interestingly, we also observed a 6-gene subset of interferon response pathway genes that was markedly induced by combo treatment but was only minimally to moderately induced by either drug alone (Figure 4F). This suggests that the combo is specifically inducing primarily neutrophil differentiation, and that aspects of interferon signaling may be uniquely induced by the combo.

We then asked why LSD1 inhibition fails to induce differentiation on its own. LSD1 is arguably the most consistently shown epigenetic regulator to drive the AML differentiation block, with numerous studies showing that its loss promotes varying degrees of myeloid differentiation. If pro-myeloid differentiation is induced by LSD1i, then why are cells falling short of terminal differentiation up on LSD1i treatment? We shed light on this question by analyzing expression trends of the direct targets of the ER-Hoxa9 transcription factor. These are genes found in our previous study^17^ to be bound by ER-Hoxa9 according to ChIP-seq and quickly downregulated upon myeloid differentiation according to RNA-seq. In ER-Hoxa9 cells – and very likely other Hox-driven AMLs – this is the most critical set of genes that must be downregulated for cells to fully differentiate. As expected, this gene signature was strongly downregulated in combo-treated cells (Figure 4G). We also observed similarly strong downregulation in SNDX-5613-treated cells. Strikingly, ORY-1001 alone had no effect on this gene set (Figure 4G). This suggests that, in this model at least, the reason LSD1 inhibitors fail to induce terminal differentiation may be that they fail to silence genes driving stemness and proliferation. Importantly, this is counter to dogma, which holds that gain of differentiation and loss of proliferation/stemness are deeply connected and can not be decoupled.

We examined the list of ER-Hoxa9 direct targets that ORY-1001 failed to downregulate, and considered these genes in light of published literature on Hox-driven AML biology. This led us to *Meis1* and *Pbx3*, which are both critical Hoxa9 heterodimerization co-factors. Both genes were strongly downregulated upon monoagent SNDX-5613 treatment and combo treatment but unchanged upon monoagent ORY-1001 treatment (Figure 4H). *Meis1*, in particular, has been proposed in multiple studies to be potentially the most important transcriptional target of the Menin/MLL complex^16,18^. If this hypothesis is correct, we would expect that loss of *Meis1* would phenocopy or near-phenocopy SNDX-5613 treatment. To test this hypothesis, we knocked out *Meis1* with CRISPR sgRNAs and tested the effect on the differentiation of ER-Hoxa9 cells both on its own and in combination with ORY-1001 treatment. In line with predictions, *Meis1* KO largely phenocopied SNDX-5613 treatment. On its own, KO had minimal effect on differentiation, as did ORY-1001 treatment. However, when combined, *Meis1* KO strongly potentiated ORY-1001, leading to high induction of the ER-Hoxa9 GFP differentiation reporter (Figure 4I).

We next interrogated the chromatin basis of the effects of combo treatment. The main prediction of our model is that LSD1 and Menin should be occupying and regulating different sets of genes; LSD1 repressing pro-differentiation genes, and Menin activating pro-stemness/proliferation genes. To test this hypothesis, we leveraged published LSD1 and GFI1B ChIP-seq datasets deriving from THP1 cells^9^. To this we added our own Menin ChIP-seq data from the closely related MOLM13 cell line. GFI1B was included because it has been proposed in multiple studies to form a key repressive complex with LSD1 in AML cells^9,19^. We first evaluated the quality of the ChIP-seq datasets by performing motif enrichment analyses. In LSD1 ChIP-seq peaks, key myeloid regulator motifs were strongly enriched, with CEBP, PU.1, Elf4, and ETS1 all scoring at p < 1e-500 (Figure 5A). Importantly, the Gfi1b motif also scored among the top motifs at p = 1e-553. This latter result strongly suggests biological accuracy of the LSD1 ChIP-seq data, as it recapitulates the known critical association between LSD1 and GFI1b. In GFI1B ChIP-seq, the top motif, at p = 1e-741, was Gfi1b itself, confirming ChIP quality (Figure 5A). Menin ChIP-seq was not as robust, but the top motif was the MLL biology-associated master transcription factor MYB (Figure 5A), and among the top Menin peaks in magnitude were the key known Menin direct targets *Meis1* and *Pbx3* (Figure 5B). These results suggest that the LSD1, GFI1B, and Menin ChIP-seq datasets are of high enough quality for further analyses.

**Figure 5.**
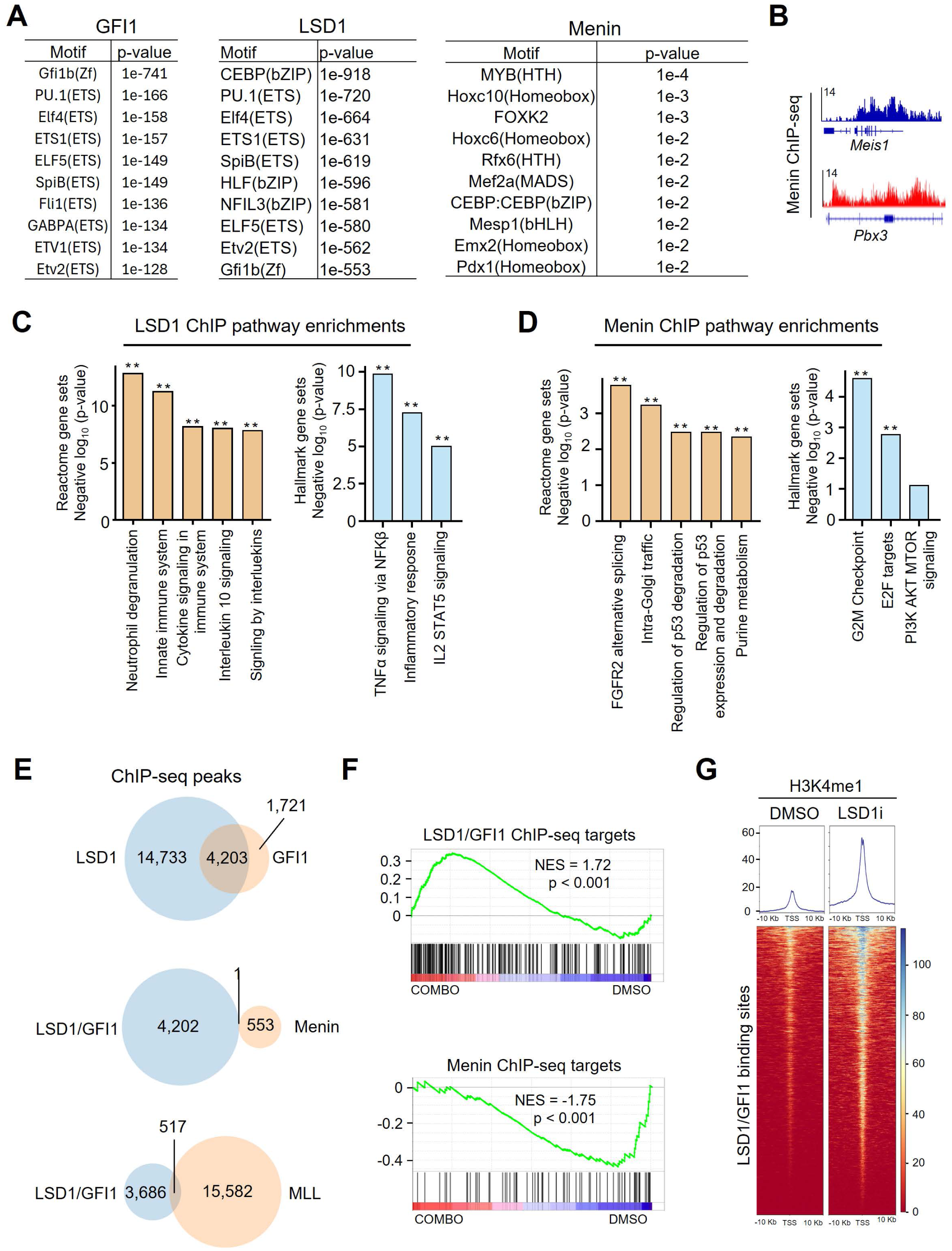
Chromatin localization of LSD1 and Menin. **A.** Motif enrichment data in peak sets from ChIP-seq of GFI1 and LSD1 in THP1 cells, and Menin in MOLM13 cells. **B.** ChIP-seq tracks showing Menin binding patterns at the *Meis1* and *Pbx3* loci. **C.** Reactome and Hallmark database pathway enrichments in list of genes with proximal LSD1 ChIP-seq peaks. **D.** Reactome and Hallmark database pathway enrichments in list of genes with proximal Menin ChIP-seq peaks. **E.** ChIP-seq heat maps of H3K4me1 signal at LSD1/GFI1 co-bound sites in cells treated with or without the LSD1 inhibitor GSK-LSD1. **F.** GSEA plots showing gene expression patterns of genes with LSD1/GFI1 co-bound or Menin-bound proximal binding sites in RNA-seq data from ER-Hoxa9 treated with ORY-1001+SNDX-5613 drug combination. **G.** Overlap of selected ChIP-seq peak lists in THP1 cells (LSD1, GFI1, and MLL) and MOLM13 cells (Menin).

We investigated which gene ontologies were enriched in LSD1 and Menin ChIP-seq datasets. The top Reactome gene set enriched in the top 10% of LSD1 ChIP-seq peaks was, as with the RNA-seq synergy signature, Neutrophil degranulation (Figure 5C). (The top 10% was used to reduce the >18,000 peak dataset to a reasonable number to be used for enrichment analyses.) The top Hallmark gene sets were TNFα signaling, Inflammatory response, and IL2 STAT5 signaling. In Menin ChIP-seq peaks, top Reactome pathways included p53 regulation and degradation, and purine metabolism, while the top Hallmark pathways were G2M checkpoint and E2F targets (Figure 5D). These findings align with our hypothesis that LSD1 is regulating pro-differentiation genes, while Menin may be regulating proliferation, metabolism, and p53-related genes.

We next asked the critical question of how much overlap there was in these ChIP-seq peak lists. We first overlapped LSD1 peaks with GFI1B peaks and found a very high degree of overlap, with 4,203 of the 5,924 GFI1B peaks overlapping an LSD1 peak (Figure 5E). Moving forward, we used this overlapping peak dataset, as it likely contains the most high-confidence LSD1 peaks. We next examined overlap between LSD1/GFIB peaks and Menin peaks. Strikingly, of the 4,203 LSD1/GFI1B peaks and the 554 Menin peaks, only 1 was overlapping. To test this in a second manner, we obtained MLL ChIP-seq data also produced in THP1 cells^20^. We examined overlap in the LSDI/GFI1B peaks with MLL peaks and again we found minimal overlap, with only 517 of the 16,089 MLL peaks overlapping the 4,203 LSD1/GFI1B peaks (Figure 5E). Taken together, these data indicate that LSD1/GFI1B and Menin/MLL are for the most part occupying different areas of the genome.

We next examined the transcriptional fate of genes bound by LSD1/GFI1B and Menin. If LSD1 is repressing genes driving myeloid differentiation, we would expect these genes to be upregulated upon combo treatment. Conversely, if Menin is activating genes driving proliferation and stemness, we would expect these genes to be downregulated upon combo treatment. Indeed, GSEA revealed precisely these trends using the full Menin target peak list and the top 10% of LSD1/GFI1B-bound genes (Figure 5F). Interestingly, Menin targets were most strongly downregulated after 24 hours (rather than 96 hours) of combo treatment, while LSD1/GFI1B targets were most strongly upregulated after 96 hours of combo treatment.

While LSD1 is an H3K4me1/2 demethylase, it is unclear whether its catalytic activity is relevant for its role in repressing myeloid differentiation genes^9^. To shed light on this question, we examined datasets^13^ containing H3K4me1 levels at LSD1/GF1B-bound loci with and without treatment with the LSD1 inhibitor GSK-LSD1 in THP1 cells. Notably, we found a dramatic increase in H3K4me1 at LSD1/GF1B-bound loci upon inhibition of LSD1 (Figure 5G). This is only correlative evidence, but would be consistent with LSD1 repressing pro-myeloid differentiation genes at least in part via H3K4me2 demethylation.

Finally, we asked whether combinatorial inhibition of LSD1 and Menin would have anti-leukemic activity in AML mouse models. To this end, we employed the commonly used MLL-AF9 retroviral overexpression syngeneic transplant model. We transplanted 500,000 MLL-AF9 cells into sublethally-irradiated C57BL/6 mice, and after confirming successful engraftment on day 10 we started treatment with the ORY-1001 + SNDX-5613 combination or vehicle control. ORY-1001 was delivered via oral gavage at 0.03 mg/kg three days per week, and SNDX-5613 was delivered at 0.1% in chow supplied by Syndax pharmaceuticals. Excitingly, we found that the drug combo dramatically extended lifespan (p = 3e-5), with several mice being cured (Figure 6A). This indicates that the ORY-1001+SNDX-5613 drug combination may be an effective treatment strategy for non-APL AML.

**Figure 6.**
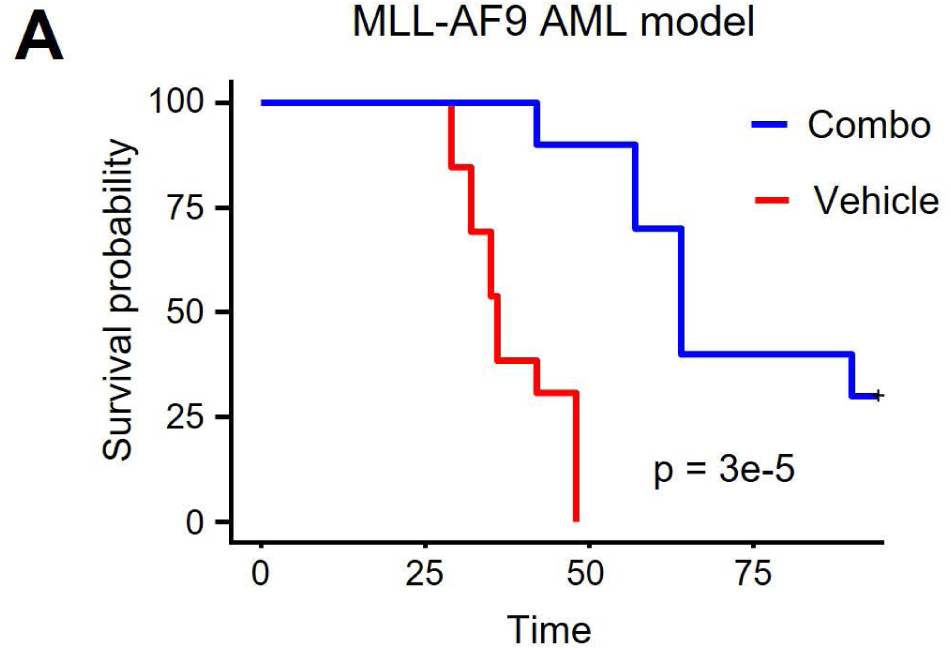
Survival of mice treated with drug combination. **A.** Kaplan-Meier plot showing survival of mice transplanted with mouse MLL-AF9 cells and treated with ORY-1001+SNDX-5613 drug combination or vehicle control.

## Discussion

Differentiation therapy of acute promyelocytic leukemia (APL) represents a landmark success in oncology^3^. Prior to differentiation therapy with ATRA and ATO, APL had the worst prognosis of all AML subtypes; now it has the best. For many years, it was thought that differentiation therapy of APL was a special case and not generalizable to non-APL AMLs. However, recent work questions this stance. IDH1/2 inhibitors, for example, have had success in the clinic and act primarily via differentiation induction for *IDH1/2-*mutant AML^7^. This suggests that differentiation therapy may be achievable in non-APL if the most critical drivers of differentiation arrest can be identified and targeted.

Here, we used differentiation-specific CRISPR screening with chromatin-focused sgRNA libraries to discover that Menin inhibition, when combined with LSD1 inhibition, synergistically induces powerful differentiation responses in non-APL AML cells. Our findings, based on cell lines, patient samples, and AML mouse models, suggest that LSD1 inhibition on its own de-represses pro-myeloid differentiation expression programs but fails to downregulate drivers of stemness and proliferation. Conversely, in certain cellular contexts at least, inhibition of Menin on its own silences drivers of stemness and proliferation but does not lead to full de-repression of myeloid differentiation genes. We propose that the combination of LSD1 and Menin inhibition thus succeeds because it directly acts on both the stemness/proliferation and the pro-differentiation gene expression modules. Importantly, these findings run counter to common models of myeloid differentiation, which hold that stemness/proliferation and pro-differentiation gene expression modules are indelibly coupled, and that one cannot be manipulated without changing the other.

The potentially most surprising aspect of our findings is that the drug combination is effective in selected MLL-WT/NPM1-WT AML models. Menin inhibitors act by displacing Menin from its interaction the MLL protein, thereby crippling the activity of the MLL complex^16^. As MLL-rearranged AMLs are driven by the high activity of the MLL fusion protein, Menin inhibitors are thought to be most effective in MLL-r AML. Additionally, it has been suggested that Menin inhibitors may also be effective in NPM1-mutant AML^21,22^. This is thought to be due to the reliance of NPM1-mutant AML on the wild type MLL protein to help drive expression of Hox-like gene expression programs, which is a critical activity of MLL in healthy myeloid cells. However, it is not thought the Menin inhibitors would be effective outside of MLL-r and NPM1-mutant AMLs, and indeed our findings are in line with this when Menin inhibitors are used as monoagents. But, very surprisingly, we find that MLL-WT/NPM1-WT AML cell lines and patient samples are sensitive to Menin inhibitors when used in combination with LSD1 inhibitors. This suggests that the MLL protein may have unappreciated roles in these cellular contexts. It will be of interest to dissect the role of the MLL complex in AMLs that are MLL-WT/NPM1-WT yet still sensitive to the LSD1i/NPM1i drug combination.

In summary, we find that LSD1 inhibitors synergize with Menin inhibitors to induce therapeutic differentiation in non-APL AML cells of a variety of genetic backgrounds. As both inhibitors are already in clinical trials as monoagents, these findings suggest the possibility that they could also be tested as a combination therapy in the clinic.

## Acknowledgments

We would like to thank Martin Carroll for helpful conversations and feedback on our findings. This work was supported by: NIH/NCI R01 CA279317 (M.A.B.) and 1F31CA281117-01 (M.F.C.R).

## Author contributions

M.A.B.: Conceptualization, experimental investigation, data analysis, manuscript writing

M.F.C.R.: Experimental investigation, data analysis, manuscript writing

J.R.: Experimental investigation, data analysis

M.V.: Experimental investigation, data analysis

F.Y.: Experimental investigation, data analysis

N.A.L.: Experimental investigation

R.P.: Experimental investigation

S.H.: Experimental investigation

K.B.: Conceptualization, data analysis

G.M.: Experimental investigation

A.K.: Experimental investigation

## Materials and Methods

### Cell culture

Human AML cell lines (U937, MOLM-13, MV4;11, NOMO-1, HL-60, K562, OCI-AML3, and THP1) were cultured in RPMI (Corning, 10-041-CV) with 10% FBS (Life Technologies, 16000044) and 1% Penicillin Streptomycin (Gibco, 15-140-122). Mouse ER-HOXA9 and Cas9-ER-HOXB8 cells were cultured in RPMI (Corning, 10-041-CV) with 10% FBS (Life Technologies, 16000044), 1% Penicillin Streptomycin (Gibco, 15-140-122), 1% - 2% stem cell factor (SCF) conditioned media (generated from a Chinese hamster ovary (CHO) cell line that stably secretes SCF), and 0.5 µM β-estradiol (Fisher Scientific MP021016562). Mouse MLL-AF9 cells (a gift from Dr. Francois Mercier at McGill University) were cultured in RPMI (Corning, 10-041-CV) with 10% FBS (Life Technologies, 16000044), 1% Penicillin Streptomycin (Gibco, 15-140-122), 1% - 2% stem cell factor (SCF) conditioned media (generated from a Chinese hamster ovary (CHO) cell line that stably secretes SCF), and 10 ng/mL IL-3 (Stemcell Technologies, 78042.1).

All cell lines were tested for mycoplasma contamination (and all confirmed mycoplasma negative) routinely, including upon each unfreeze and then at least monthly during periods of continued culture. Testing was performed using the Universal Mycoplasma Detection Kit (ATCC, 30-1012K) according to the manufacturer’s instructions.

Bone marrow or peripheral blood mononuclear cells from patients with AML (AML_2263 and AML_6740) were obtained from the Stem Cell and Xenograft Core (SCXC) facility at the University of Pennsylvania. Human primary AML cells were cultured in IMDM (Gibco, 12440-053) with 2% FBS (Life Technologies, 16000044) and 1% Penicillin Streptomycin (Gibco, 15-140-122) at a concentration of 2 million cells/mL.

### Virus packaging and transduction

For lentivirus packaging, the target vector and pCMV-VSV-G and pCMV-dR8.2 dvpr lentiviral packaging plasmids (gifts from Dr. Bob Weinberg at the Massachusetts Institute for Technology), Addgene plasmid # 8454 and Addgene plasmid # 8455) were co-transfected into 293T cells (Clontech, 632180) using PEI reagent (Polysciences, 23966-1). Lentiviral particles were collected 48 and 72 hours after transfection, filtered and added to target cells with 8µg/mL polybrene (Sigma, TR-1003-G). For retrovirus packaging, the target vector was transfected into the GPG29 packaging cell line using Lipofectamine 2000 (Invitrogen). Retroviral particles were collected 72-96 hours after transfection, filtered and added to target cells with 8µg/mL polybrene (Sigma, TR-1003-G). Transduced cells were then selected by appropriate antibiotics or cell sorting 48 hours after infection.

### Colony formation assays

Clonogenic potential was assessed by seeding the indicated number of cells in methylcellulose media. To make 100mL complete methylcellulose media, 40mL methylcellulose base media (Stem Cell Technologies, MethoCult H4100) was supplemented with IMDM (Gibco, 12440-053), 10% FBS (Life Technologies, 16000044), 1% Penicillin Streptomycin (Gibco, 15-140-122) and other desired supplements as described previously in cell culture section. The complete media was then used to resuspend the cells. Number of colonies was counted two weeks following seeding.

### Animal experiments

500,000 mouse MLL-AF9 cells were transplanted by tail vein injection into C57BL/6 mice 24 hours after irradiating the whole body of mice with a single dose of 4.5 Gy. On day 10 treatment with ORY-1001 and SNDX-5613 was initiated. ORY-1001 was delivered at 0.03 mg/kg via oral gavage 3X/week. SNDX-5613 was delivered at 0.1% weight x weight in mouse chow. Kaplan-Meier survival curves and log-ranks tests were performed using the Survival R package.

### ChIP-seq and analysis

5-10 million cells were used for ChIP of Menin. Cells were washed with PBS, crosslinked with 1% formaldehyde for 10 minutes at room temperature, and then quenched with 125 mM glycine for 5 minutes. The isolated nuclei were resuspended in 1 mL nuclei lysis/sonication buffer and sonicated with Covaris S220 sonicator with following parameters: Peak Intensity – 140, Duty Factor – 5, Cycles per Burst – 200, Time – 60 seconds on, 30 seconds off for 16 cycles. After centrifugation at 4 degrees with 13,500 rpm for 10 min, soluble chromatin was then used to perform immunoprecipitation, while 5% sample kept as input DNA. Immunoprecipitation was performed with 5 – 10 μg Menin antibody overnight at 4 degrees with rotation. Products were then incubated with magnetic Protein G Dynabeads and then washed sequentially using low-salt, high-salt, LiCl buffer, and TE buffer. Bound DNA was then eluted, reverse-crosslinked, incubated with RNase A and Proteinase K. DNA samples were purified using QIAquick PCR Purification Kit (Qiagen) and used for preparation of ChIP-seq libraries. ChIP-seq libraries were prepared following the NEBNext Ultra II DNA Library Prep Kit (New England Biolabs) protocol, using NEBNext Multiplex Oligos Index Primers Sets. Libraries were pooled and sequenced on the NextSeq500/550 (Illumina, 75 cycles High Output kit v2.0) to generate 75 bp single end reads or paired end reads. Sequencing reads were aligned to the human (hg38) using Bowtie2 with default settings^23^. Resulting sam files were filtered for uniquely aligned reads, converted to bam files, sorted, and marked for duplicated reads using SAMtools^24^. Peaks calling was performed using MACS2. Peak overlaps were analyzed with bedtools, and heat maps were generated with deepTools. Homer was used for motif enrichment in peak sets, and EnrichR was used to identify gene ontologies enriched in peak sets. LSD1 ChIP-seq and GFI1B ChIP-seq data were obtained from Maiques-Diaz *et al.*^9^, MLL ChIP-seq data were obtained from Prange *et al.*^20^, and H3K4me1 ChIP-seq data were obtained from Hosseini *et al*.^13^

### RNA purification and RNA-seq analysis

RNA samples of 3 biological replicates per treatment were extracted from cultured cells using Qiagen RNeasy Kits following manufacturer’s instructions. RNA was then sent out for library preparation and next-generation sequencing to a commercial company, Novogene (California, USA). Raw counts of gene transcripts were derived from raw fastq files using the alignment-independent quantification tool, Salmon (https://combine-lab.github.io/salmon/), with standard settings. The raw count matrix was then imported into R-studio and utilized as input for DESeq2 analysis following the vignette of the package for normalization, differential gene expression analysis, and unbiased clustering analysis, including principal component analysis^25^. The output of DESeq2 was used as the input for pre-ranked based GSEA for enrichment of functional pathways and gene signatures (https://www.gsea-msigdb.org/gsea/index.jsp). Detailed Scripts and parameters used for each step of analysis could be provided by request to the authors.

### Pooled CRISPR-Cas9 screen and screen analysis

The human and mouse epi-RBP CRISPR sgRNA libraries were constructed using methodology described in detail in recent publications^26,27^. Briefly, the sgRNA oligos were designed according to a published method^28^, and 100 non-targeting control sgRNAs were also included. The sgRNA library was synthesized using array synthesis by CustomArray, Inc and cloned as a pool into the lentiGuide-puro transfer plasmid via Gibson ligation reaction (NEB). After ligation, the library was transformed into electrocompetent cells (Lucigen) for amplification.

For CRISPR screens, Cas9-ER-Hoxb8 or Cas9-U937 cells were first transduced with the mouse (for Cas9-ER-Hoxb8) or human (for Cas9-U937) sgRNA lentivirus library at an MOI of 0.2. At least 200x coverage of the sgRNA library was maintained throughout the screens. 24 hours after infection cells were selected with puromycin, and 72 hours after infection cells were split into one branch that received DMSO and another branch that received 10 nM ORY-1001. One week later, AML cells were sorted via FACS based on surface expression of CD11b into the top 2% CD11b-high and the bottom 2% CD11b-low samples for library preparation. Genomic DNA from sorted and bulk AML cells was harvested using the QIAamp DNA mini kit (Qiagen). Sequencing libraries were then prepared following previously described protocols^29^. All prepared genomic DNA was used for library preparation for maintaining library coverage. PCR amplified library samples were purified with the QIAquick PCR purification kit (Qiagen) followed by gel extraction with the QIAquick gel extraction kit (Qiagen). The barcoded libraries were then pooled at an equal molar ratio and sequenced on a NextSeq500/550 (Illumina, 150 cycles High Output kit v2.0) to generate 150 bp single end reads. MAGeCK software was used for screen analysis^30^. Briefly, the resulted sequencing data were de-barcoded, merged, and the 20 bp sgRNA sequence was aligned to the reference sgRNA library without allowing for any mismatches. The read counts were calculated for each sgRNA using the method normalizing to the non-targeting sgRNAs. Differential analysis of sgRNA and targeted genes was also done following the MAGeCK instructions with standard parameters. Detailed Scripts and parameters used for each step of analysis could be provided by request to the authors.

### Mouse models

C57BL/6J mice used in this study were 8-week female mice purchased from The Jackson Laboratory (000664) and maintained in the mouse facility at the School of Veterinary Medicine at the University of Pennsylvania. All mouse procedure protocols utilized in study this were in accordance with, and with the approval of, the Institutional Animal Care and Use committee (IACUC). All mouse experiment procedures used in this study were performed following the National Institutes of Health guidelines.

### Colony formation assay

Clonogenic potential was assessed by seeding the indicated number of cells in methylcellulose media. For cell lines: to make 100mL complete methylcellulose media, 40mL methylcellulose base media (Stem Cell Technologies, MethoCult H4100) was supplemented with IMDM (Gibco, 12440-053), 10% FBS (Life Technologies, 16000044), 1% Penicillin Streptomycin (Gibco, 15-140-122) and other desired supplements as described previously in cell culture section. The complete media was then used to resuspend the cells. Number of colonies was counted two weeks following seeding.

### Flow cytometry and cell sorting

For flow cytometric analyses of cell surface CD11b, cells were washed with PBS then stained with PE-CD11b antibody (Biolegend, #101207) at room temperature for 15 minutes and then washed twice with cold PBS for flow cytometric analysis. Flow cytometry was performed on BD LSRFortessa (BD Biosciences) and analyzed with FlowJo software (Treestar). Fluorescence activated cell sorting (FACS) of AML cells was performed by MoFlo Astrios (Beckman) or BD Jazz (BD Biosciences) according to the manufacturer’s instructions.

### In vitro drug treatment

Cells were treated in triplicates and treated with indicated drugs or DMSO. Chemicals are listed below:

ORY-1001, Cayman, 19136

SNDX-5613, MedChemExpress, HY-136175

GSK-LSD1, Sellekchem, S7574

### Superoxide Anion Asssay

Superoxide Anion assay were performed using Sigma #CS1000 kit following product instructions. Luminescence was measured using the EnVision (PerkinElmer) plate reader every 10 minutes in 4 hours.

### Software and Statistical analysis

PRISM software and R were used for data processing, statistical analysis, and result visualization (http://www.graphpad.com). The R language and environment for graphics (https://www.r-project.org) was used in this study for the bioinformatics analysis of CRISPR screen, RNA-seq, and ChIP-seq data. The R packages used for all analysis described in this manuscript were from the Bioconductor and CRAN. On graphs, bars represent either standard deviation (SD) or standard error of mean (SEM), as indicated in legends. For all figures, p<0.05 was considered statistically significant, *indicates p<0.01, and **p<0.001.

## Supplemental Figure Legends

**Figure S1.**
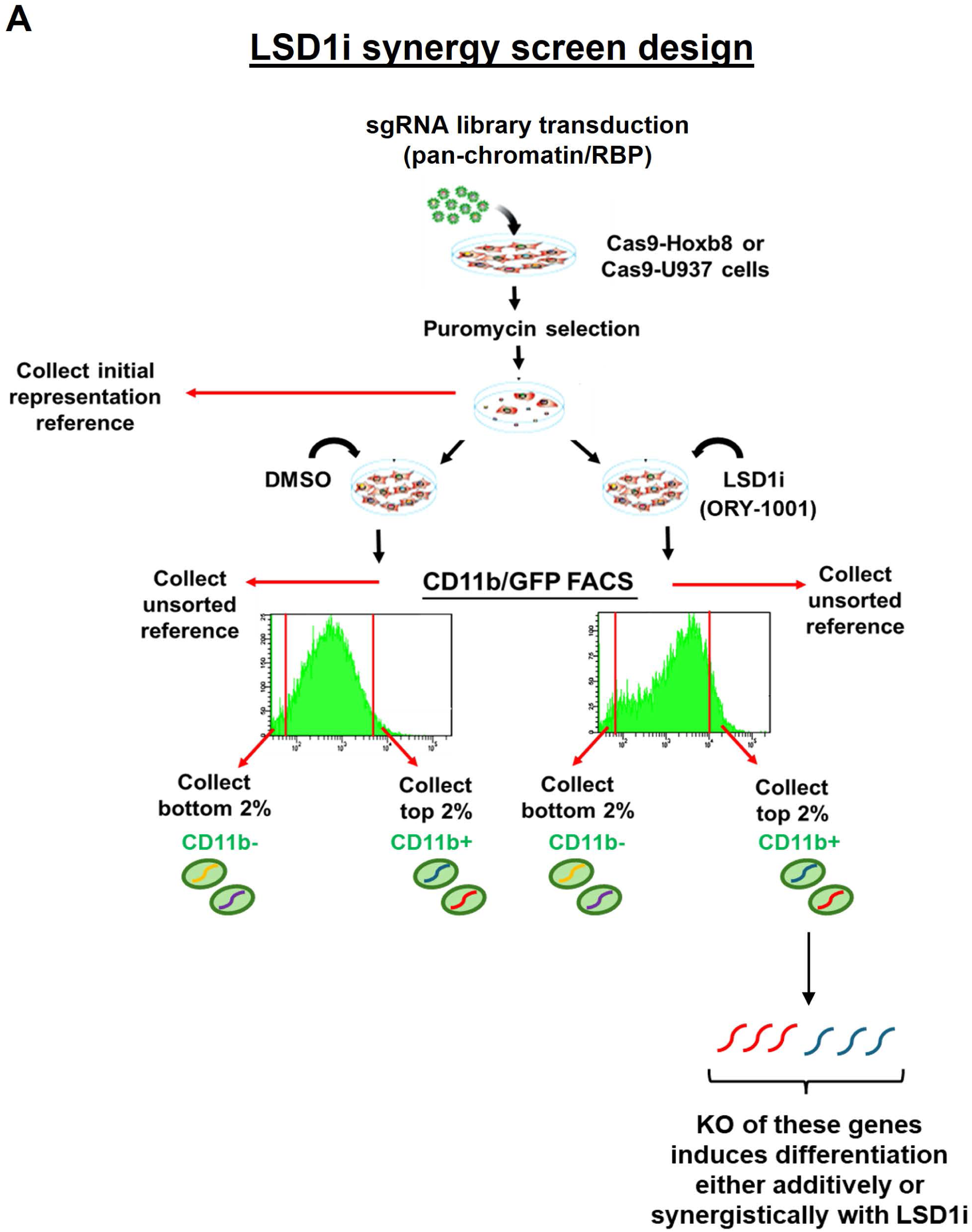
LSD1i synergy screen design. **A.** Schematic showing scheme for differentiation-specific chromatin-focused screen for genes whose inhibition synergizes with ORY-1001 to induce differentiation as gauged by CD11b as a differentiation readout.

**Figure S2.**
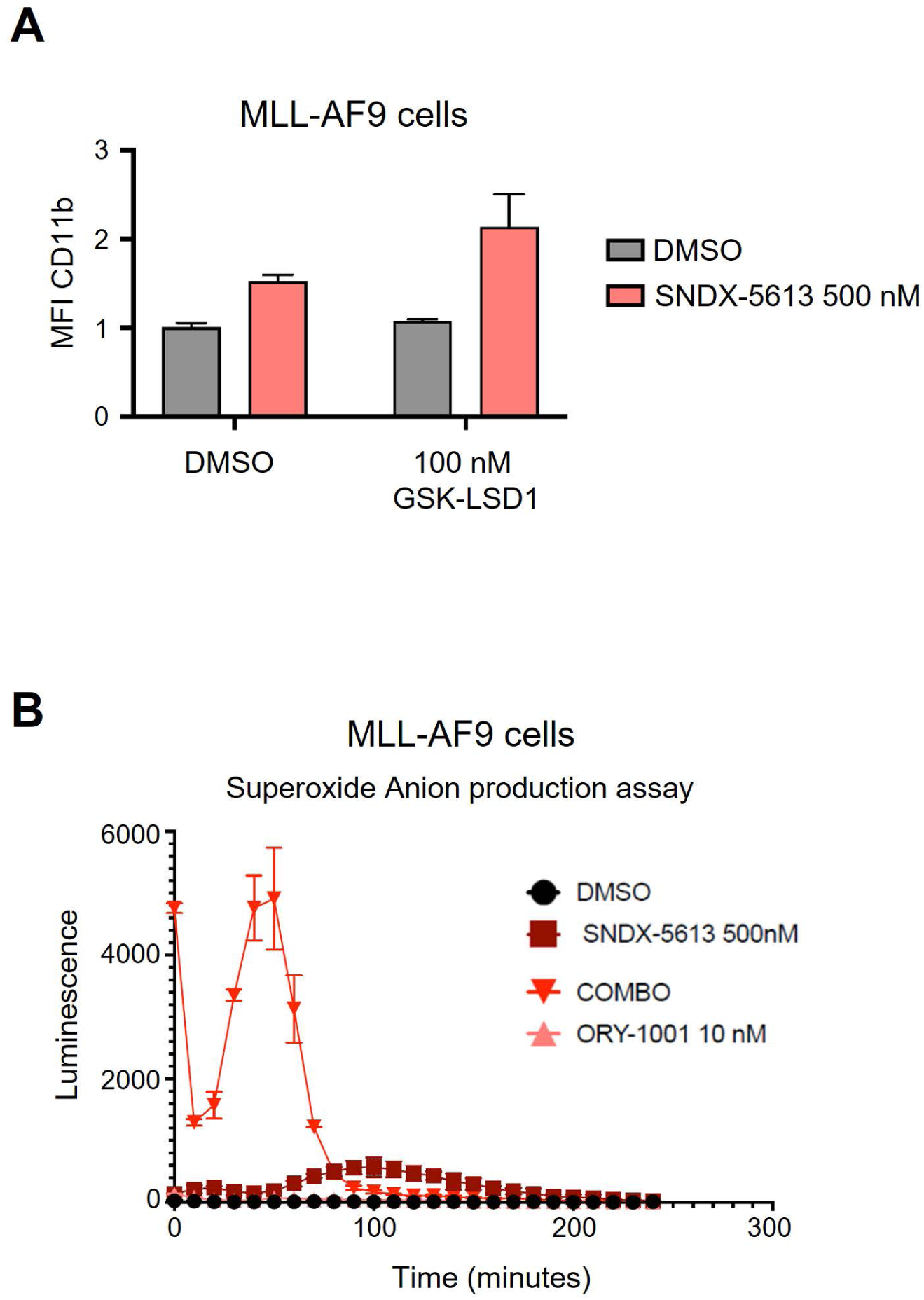
LSD1 inhibition synergizes with Menin inhibition to induce differentiation. **A.** CD11b differentiation response assays in MLL-AF9 cells treated with GSK-LSD1, SNDX-5613, the combination, or neither. **B.** Superoxide anion response of MLL-AF9 cells upon treatment with ORY-1001, SNDX-5613, the combination, or neither.

## References

1. Sung, H. et al. Global Cancer Statistics 2020: GLOBOCAN Estimates of Incidence and Mortality Worldwide for 36 Cancers in 185 Countries. 10.3322/caac.21660 doi:10.3322/caac.21660.

2. Wertheim, G. B. W., Hexner, E. & Bagg, A. Molecular-Based Classification of Acute Myeloid Leukemia and Its Role in Directing Rational Therapy. Mol Diagn Ther 16, 357–369 (2012).

3. Tenen, D. G. Disruption of differentiation in human cancer: AML shows the way. Nat Rev Cancer 3, 89–101 (2003).

4. Huang, M. E. et al. All-trans retinoic acid with or without low dose cytosine arabinoside in acute promyelocytic leukemia. Report of 6 cases. Chin Med J (Engl) 100, 949–953 (1987).

5. Petrie, K., Zelent, A. & Waxman, S. Differentiation therapy of acute myeloid leukemia: past, present and future. Curr Opin Hematol 16, 84–91 (2009).

6. Goldman, S. L. et al. Epigenetic Modifications in Acute Myeloid Leukemia: Prognosis, Treatment, and Heterogeneity. Front Genet 10, 133 (2019).

7. Yen, K. et al. AG-221, a First-in-Class Therapy Targeting Acute Myeloid Leukemia Harboring Oncogenic IDH2 Mutations. Cancer Discov 7, 478–493 (2017).

8. Schenk, T. et al. Inhibition of the LSD1 (KDM1A) demethylase reactivates the all-trans-retinoic acid differentiation pathway in acute myeloid leukemia. Nat Med 18, 605–611 (2012).

9. Maiques-Diaz, A. et al. Enhancer Activation by Pharmacologic Displacement of LSD1 from GFI1 Induces Differentiation in Acute Myeloid Leukemia. Cell Rep 22, 3641–3659 (2018).

10. The Histone Demethylase KDM1A Sustains the Oncogenic Potential of MLL-AF9 Leukemia Stem Cells. Cancer Cell 21, 473–487 (2012).

11. Fang, Y., Liao, G. & Yu, B. LSD1/KDM1A inhibitors in clinical trials: advances and prospects. J Hematol Oncol 12, 1–14 (2019).

12. Duy, C. et al. Rational targeting of cooperating layers of the epigenome yields enhanced therapeutic efficacy against AML. Cancer discovery 9, 872 (2019).

13. Hosseini, A. et al. Perturbing LSD1 and WNT rewires transcription to synergistically induce AML differentiation. Nature 642, 508–518 (2025).

14. Wang, G. G. et al. Quantitative production of macrophages or neutrophils ex vivo using conditional Hoxb8. Nature Methods 3, 287–293 (2006).

15. Sykes, D. B. et al. Inhibition of Dihydroorotate Dehydrogenase Overcomes Differentiation Blockade in Acute Myeloid Leukemia. Cell 167, 171–186.e15 (2016).

16. Krivtsov, A. V. et al. A Menin-MLL Inhibitor Induces Specific Chromatin Changes and Eradicates Disease in Models of MLL-Rearranged Leukemia. Cancer Cell 36, 660–673.e11 (2019).

17. Blanco, M. A. et al. Chromatin-state barriers enforce an irreversible mammalian cell fate decision. Cell Rep 37, 109967 (2021).

18. Heikamp, E. B. et al. The menin-MLL1 interaction is a molecular dependency in NUP98-rearranged AML. Blood 139, 894–906 (2022).

19. Venhuizen, J. et al. GFI1B and LSD1 repress myeloid traits during megakaryocyte differentiation. Commun Biol 7, 1–9 (2024).

20. Prange, K. H. M. et al. MLL-AF9 and MLL-AF4 oncofusion proteins bind a distinct enhancer repertoire and target the RUNX1 program in 11q23 acute myeloid leukemia. Oncogene 36, 3346– 3356 (2017).

21. Uckelmann, H. J. et al. Mutant NPM1 directly regulates oncogenic transcription in acute myeloid leukemia. Cancer Discov 13, 746–765 (2023).

22. Xqd, W. et al. Mutant NPM1 Hijacks Transcriptional Hubs to Maintain Pathogenic Gene Programs in Acute Myeloid Leukemia. Cancer discovery 13, (2023).

23. Langmead, B. & Salzberg, S. L. Fast gapped-read alignment with Bowtie 2. Nat Methods 9, 357– 359 (2012).

24. Li, H. et al. The Sequence Alignment/Map format and SAMtools. Bioinformatics 25, 2078 (2009).

25. Love, M. I., Huber, W. & Anders, S. Moderated estimation of fold change and dispersion for RNA-seq data with DESeq2. Genome Biol 15, 550 (2014).

26. Shi, Z. et al. Mettl17, a regulator of mitochondrial ribosomal RNA modifications, is required for the translation of mitochondrial coding genes. 10.1096/fj.201901331R doi:10.1096/fj.201901331R.

27. Yan, F. et al. KAT6A and ENL Form an Epigenetic Transcriptional Control Module to Drive Critical Leukemogenic Gene-Expression Programs. Cancer Discov 12, 792–811 (2022).

28. Chen, S. et al. Genome-wide CRISPR screen in a mouse model of tumor growth and metastasis. Cell 160, 1246–1260 (2015).

29. Joung, J. et al. Genome-scale CRISPR-Cas9 Knockout and Transcriptional Activation Screening. Nat Protoc 12, 828–863 (2017).

30. Li, W. et al. MAGeCK enables robust identification of essential genes from genome-scale CRISPR/Cas9 knockout screens. Genome Biol 15, 554 (2014).

